# Emergence of a cephalosporin reduced susceptible Neisseria gonorrhoeae clone between 2014-2019 in Amsterdam, the Netherlands, revealed by a genomic population analysis

**DOI:** 10.1101/2020.11.02.364463

**Authors:** Jolinda de Korne-Elenbaas, Sylvia M. Bruisten, Henry J.C. de Vries, Alje P. van Dam

**Author notes:** Corresponding author: Jolinda de Korne-Elenbaas.

## Abstract

**Background:** Emerging resistance to cephalosporins in *Neisseria gonorrhoeae (Ng)* is a major public health threat, since these are considered antibiotics of last resort. Continuous surveillance is needed to monitor the circulation of reduced susceptible and resistant strains.

**Aim:** For the purpose of epidemiological surveillance, a genomic population analysis was performed on *Ng* isolates from Amsterdam with a focus on ceftriaxone reduced susceptible isolates.

**Methods:** Whole genome sequences were obtained from 318 isolates from 2014-2019 from Amsterdam, the Netherlands. Isolates were typed according to the Multi-Locus Sequence Typing (MLST) scheme and resistance markers were identified. Phylogenetic trees were created to identify genetic clusters and to compare Dutch- and global MLST7827 isolates.

**Results:** The majority of ceftriaxone reduced susceptible isolates belonged to MLST7363, MLST1901 and MLST7827. MLST7363 and MLST1901 were dominant reduced susceptible strains from 2014-2016 although MLST7827 emerged and became dominant during 2017-2019. MLST7827 reduced susceptible isolates carried a non-mosaic *penA* gene with A501V mutation and *porB* G121K/A121D mutations, which were lacking in susceptible MLST7827 isolates. Phylogenetic analysis of all publicly available MLST7827 isolates showed strong genetic clustering of Dutch- and other European MLST7827 isolates.

**Conclusion:** Ceftriaxone reduced susceptible MLST7827 isolates emerged during recent years in Amsterdam. Co-occurrence of *penA* A501V and *porB* G121K/A121D mutations was strongly associated with ceftriaxone reduced susceptibility. Genetic clustering of Dutch- and other European MLST7827 isolates indicates extensive circulation of this strain in Europe. Therefore, close monitoring of the spread of this strain with an alarming susceptibility profile is needed.

## Introduction

Infection with *Neisseria gonorrhoeae (Ng)* causes the sexually transmitted infection (STI) gonorrhoea, which currently poses a major public health threat due to the emergence of multidrug resistant strains. The World Health Organization and the European Centre for Disease Prevention and Control have set up Gonococcal Antimicrobial Surveillance Programs (WHO- or Euro-GASP) to monitor susceptibility patterns and track emergence of resistant strains on global or European level (1)(2). These international surveillance initiatives provide emerging resistance data necessary to update treatment guidelines. Increasing resistance of *Ng* against multiple antibiotics has led to the current treatment recommendation with the last-resort extended-spectrum cephalosporin ceftriaxone, together with azithromycin as dual therapy (3). Since the benefit of dual therapy is not evidence-based and emerging high-level azithromycin resistance has been found in many countries (4), the Dutch, French and UK treatment guidelines recommend ceftriaxone monotherapy (5). WHO guidelines endorse monotherapy as well, provided that local resistance data confirming susceptibility to ceftriaxone is available (3). However, single cases of ceftriaxone resistant isolates have been reported over the last years, underlining the need for continuous surveillance of circulating strains (6–9).

Although ceftriaxone resistant strains have only been reported in a few countries so far, a drift towards higher ceftriaxone Minimum Inhibitory Concentrations (MICs) is observed worldwide, indicating a global reduction in susceptibility (10). Ceftriaxone resistance appeared to be associated with carriage of a mosaic penicillin-binding protein A (*penA)* gene (parts of the gene are exchanged with other *Neisseria* species) or a *penA* A501 amino acid mutation (11)(12). Genotypic characterization of the ceftriaxone resistant isolates found in Japan, France and the United Kingdom showed that those isolates carried the mosaic *penA* gene and belonged to Multi-Locus Sequence Type (MLST) 1901/1903 and Neisseria Gonorrhoeae Multi-Antigen Sequence Type (NG-MAST) 1407 (6,7,9). A ceftriaxone resistant isolate from Singapore belonged to different STs but did contain the same mosaic *penA* gene (8).

Isolates belonging to MLST1901 and NG-MAST1407, and containing the mosaic *penA* were most prevalent among ceftriaxone reduced susceptible isolates as well, which suggests that tracking this strain is most important for monitoring emerging resistance. However, Osnes et al. recently reported the emergence of a reduced susceptible strain between 2016-2018 in Norway, belonging to MLST7827 and carrying a non-mosaic *penA* allele with A501V mutation (13). They showed that this strain with an alarming antimicrobial resistance profile likely originated from Asia and potentially circulates in Europe. Analysing Dutch reduced susceptible isolates from 2009-2017, De Laat et al. found a shift from a mosaic *penA* allele towards a non-mosaic *penA* allele with A501 mutation (14). In this study, the genetic change among reduced susceptible isolates in Amsterdam is further examined. Whole genome sequences (WGS) were used for genomic characterization of ceftriaxone reduced susceptible isolates and a representative part of the susceptible gonococcal population isolated from 2014-2019 in Amsterdam. We aimed to identify genomic characteristics associated with reduced susceptibility to ceftriaxone in the Amsterdam gonococcal population.

## Methods

### Isolate details and selection

The Public Health Laboratory of Amsterdam possesses a large collection of *Ng* isolates, mainly obtained from STI outpatient clinic visitors in Amsterdam. After isolation, all *Ng* isolates were inoculated on a plate with BBL GC-Lect Agar (Becton, Dickinson and Company), whereafter a pure colony was grown on a GC agar plate enriched with 1% Iso VitaleX (BioTRADING Benelux BV). Minimum Inhibitory Concentrations (MIC) for azithromycin, ciprofloxacin and ceftriaxone were determined for all isolates using E-tests according to manufacturer’s instructions (bioMérieux SA). Cultured colonies were stored at - 80°C until further use. Ciprofloxacin clinical breakpoints were determined according to EUCAST. For azithromycin, isolates with MIC<0.5 μg/mL were assigned susceptible, MIC 0.5 μg/mL intermediate and MIC≥1.0 μg/mL resistant (ECOFF). For ceftriaxone, isolates with a MIC≤0.016 μg/mL were assigned susceptible, MIC 0.016-0.094 μg/mL intermediate susceptible and MIC≥0.094 μg/mL reduced susceptible. Ceftriaxone resistant strains with MIC>0.125 μg/mL were not found during the study period.

For genomic characterization of ceftriaxone reduced susceptible strains circulating in Amsterdam, all 82 isolates from January 2014 - July 2019 with a ceftriaxone MIC≥0.094 μg/mL were selected for WGS. To characterize the total gonococcal population circulating in Amsterdam, 244 out of 7241 available isolates from the same time frame with a ceftriaxone MIC<0.094 μg/mL were selected for WGS as well. Isolates were randomly selected after stratification on year of isolation and ceftriaxone MIC: for each reduced susceptible strain, 3 isolates with MIC<0.094 μg/mL from the same year of isolation were randomly selected. Stratification on ceftriaxone MIC was done to get a distribution of MICs<0.094 μg/mL in the selection similar to the distribution of MICs<0.094 μg/mL in the total gonococcal population of Amsterdam.

### DNA isolation and whole genome sequencing

Selected isolates were grown overnight on chocolate blood agar plates and DNA was extracted from pure cultures. Isolates were sequenced on the Illumina MiSeq or Illumina NovaSeq 6000 platform (the latter was chosen for higher throughput). For Illumina MiSeq sequencing, DNA was extracted using isopropanol precipitation after lysis with NucliSENS easyMAG Lysis buffer (bioMérieux SA) enriched with glycogen (40 μg/mL). The pellet was washed twice in 70% EtOH and dissolved in 50 μL Tris-HCl at pH 8.0. DNA sequencing libraries were prepared with KAPA HTP library preparation kit (Roche Life Sciences) and Nextflex Dual-Indexed DNA barcodes (Bioo Scientific), and 300 bp paired-end sequenced. Regarding Illumina NovaSeq 6000 sequencing, DNA was extracted from harvested bacteria in DNA/RNA Shield buffer using ZymoBIOMICS™ MagBead DNA kit (ZYMO RESEARCH). DNA sequencing libraries were prepared with the Nextera XT DNA Library Preparation kit with IDT for Illumina DNA/RNA UD Indexes (Illumina) and 150 bp paired-end sequenced. All raw reads are available in the European Nucleotide Archive under accession number PRJEB40983.

### Bioinformatic analyses

Default settings were used unless noted otherwise. Raw sequence reads were trimmed and adapters were removed with fastp v0.20.0 (15). Reads were mapped to reference genome FA1090 (NC_002946.2) with minimap2 v2.17 (16) to calculate coverage using the samtools package v1.11 (17). Isolates were excluded if coverage <95%. Reads were assembled with Skesa v2.3.0 with a minimum contig length of 500 bp (18) and assembly quality was assessed with QUAST v5.0.2 (19). For isolates with a total assembly length of >2.1 Mbp Kraken2 v2.0.8 was used to check for contamination (20). Variants were called with Snippy v4.4.0 using reference genome FA1090 and a full core genome alignment was created with the snippy-core option (https://github.com/tseemann/snippy). Gubbins v2.3.4 was used to identify regions of recombination in the core alignment and to create a phylogenetic tree based on a recombination filtered variant alignment (21), by using the general time reversible model with gamma distribution (GTR-GAMMA) in RAxML v8.2.12 (22). The phylogenetic tree with metadata was visualized using iTOL and legends were added with PDF version Pro (23). Bayesian Analysis of Population Structure (BAPS) was performed using the rhierBAPS package v1.1.2 with R v3.6.3 (24).

Isolates were typed according to the Multi-Locus Sequence Typing scheme in pubMLST (25), by using mlst v2.18.0 (https://github.com/tseemann/mlst). Novel MLSTs were submitted to the pubMLST database. *Ng* multi--antigen sequence typing (NG-MAST) was done using ng-master v0.5.5 (26) and *Ng* sequence typing for antimicrobial resistance (NG-STAR) was done according to the NG-STAR database (27) using the pyngSTar.py script (https://github.com/leosanbu/pyngSTar). Novel NG-STAR STs were submitted to the NG-STAR database. Raw reads were mapped against all 23S rRNA reference sequences in the pubMLST database to identify heterogeneous A2059G and C2611T mutations in the 4 different alleles by using Ariba v2.14.4 (28). Snakemake v5.6.0 was used for workflow management (29) and the pipeline is freely available at https://github.com/jolindadekorne/Genomic-population-analysis-of-Neisseria-gonorrhoeae.

### Comparison of MLST7827 isolates

The genetic relatedness of Dutch MLST7827- and global MLST7827 isolates was assessed. PubMLST contains 224 *Ng* isolates belonging to MLST7827 (August 2020), including the Dutch isolates. Depending on availability, raw sequence reads were downloaded from NCBI or contigs from pubMLST. Raw sequence reads were trimmed and adapters were removed with fastp. One Dutch MLST7827 isolate was sequenced on both the Illumina MiSeq and the MinION Nanopore platforms. Hybrid assembly was performed using Unicycler v0.4.8 (30), yielding a circular chromosome and a circular plasmid. Variants were called with Snippy v4.6.0 using the Dutch hybrid assembly as reference genome, either using the raw reads or the contigs with the --ctgs option. A recombination corrected phylogenetic tree was created and visualized as described above. Median Single Nucleotide Polymorphisms (SNPs) distance per main genetic cluster was calculated using snp-dists v0.7.0 on the filtered variant alignment (https://github.com/tseemann/snp-dists).

### Statistical analyses

Associations between patient- and/or isolate characteristics were identified with Chi squared or Fisher’s exact test using the Bonferroni correction method in case of multiple testing. Cramer’s V association was calculated when testing for association between two categorical variables. All statistical analyses were performed in R v3.6.3.

## Results

### Sequencing data

A total of 326 *Ng* isolates were selected for WGS of which 4 were excluded due to non-viable cultures. Quality control of sequencing data and assembly led to exclusion of 3 isolates due to coverage <95% and 1 isolate due to contamination, resulting in 318 sequences usable for further analyses. Core genome alignment resulted in an alignment length of 2,153,922 sites. Recombination filtering of the variable sites in the core variant alignment resulted in a variant alignment of 18,683 sites, which was used for creation of the phylogenetic tree.

### Patient characteristics

The 318 isolates were derived from 314 patients: 8 isolates were obtained from 4 patients from two different anatomical locations. Isolates were mainly obtained from men who have sex with men (MSM) (82%) and isolated from the anus (48%). The median age of all patients was 30 years and the majority of patients were aged between 24-34 years (51%) (Table 1).

**Table 1.** Patient- and isolate characteristics.

| **Number of patients** |  | **314** |  |
| --- | --- | --- | --- |
| **Sex** | Male | 286 | (91%) |
|  | Female | 28 | (9%) |
| **Age** | Median age<sup>a</sup> | 30 | (16-65) |
|  |  |  | (18.8%) |
|  | <24 | 59 |  |
|  | 24-34 | 159 | (50.6%) |
|  | >=35 | 93 | (29.6%) |
|  | NA | 3 | (1.0%) |
| **Sexual preference** | MSM | 258 | (82.2%) |
|  | heterosexual | 41 | (13.1%) |
|  | bisexual | 12 | (3.8%) |
|  | NA | 3 | (1.0%) |
| **Number of isolates sequenced** |  | **318<sup>b</sup>** |  |
| **Year of isolation** | 2014-2016 | 128 | (40%) |
|  | 2017-2019 | 190 | (60%) |
| **Anatomical location** | Anus | 152 | (47.8%) |
|  | Urethra | 89 | (28.0%) |
|  | Vagina/cervix | 14 | (4.4%) |
|  | Tonsil | 62 | (19.5%) |
|  | Other | 1 | (0.3%) |

<sup>a</sup>Median (range)
<sup>b</sup>From 4 patients, 2 isolates were obtained from 2 anatomical locations.

### Genomic epidemiology and characterization of resistance mutations

A midpoint rooted phylogenetic tree was created based on the recombination filtered variant alignment and two separate lineages were identified (Figure 1). Isolates in lineage A (n=102) were significantly associated with females, age <24 and heterosexual intercourse. Isolates in lineage B (n=216) were mainly from patients reporting homosexual or bisexual intercourse (96%). Isolates from 2017-2019 and isolates resistant against azithromycin, ciprofloxacin or ceftriaxone were significantly overrepresented in lineage B (Table S1).

**Figure 1.**
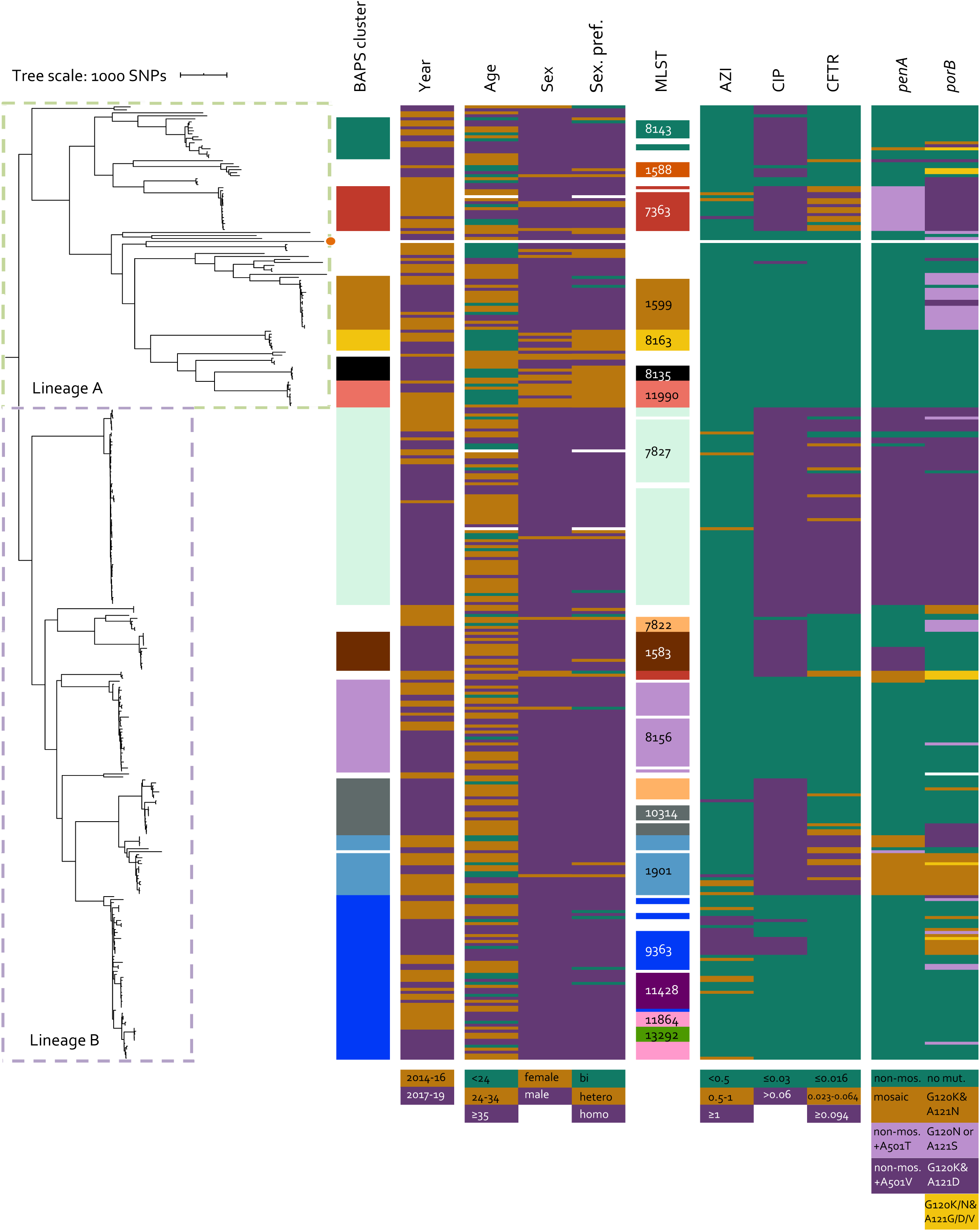
Recombination filtered midpoint-rooted phylogenetic tree based on core genome SNPs including 318 *Neisseria gonorrhoeae* isolates from 2014-2019 from Amsterdam, the Netherlands. FA1090 strain was used as reference strain and its branch is visualized with an orange dot. Metadata includes: main clusters determined with Bayesian Analysis Population Structure analysis at level 1; year of isolation; patient characteristics age, sex and sexual preference; Multi-Locus Sequence Type (MLST) clusters containing ≥ 5 isolates; Minimum Inhibitory Concentrations for azithromycin (AZI), ciprofloxacin (CIP) and ceftriaxone (CFTR); types of penA and porB genes. Phenotypic data is visualized as susceptible (green), intermediate susceptible (orange) and resistant/ reduced susceptible (purple). White bars indicate missing data. Two separate lineages are defined with dashed lined boxes.

Bayesian Analysis of Population Structure (BAPS) clustering resulted in 14 clusters of which 12 main clusters are visualized in Figure 1. MLSTs were determined for 317/318 (100%) isolates, yielding 56 different MLSTs of which 26 were found for only a single isolate. One isolate could not be assigned a MLST because of one incomplete locus. The 17 MLST clusters that contained ≥ 5 isolates were defined as main MLST clusters, with MLST7827 being the largest cluster containing 63 isolates. A strong association was found between BAPS cluster and MLST (Cramer’s V = 0.91). MLST8135, MLST8163 and MLST11990 were only found in patients reporting heterosexual intercourse and MLST11990 was significantly associated with females (Table S2). NG-MAST and NG-STAR types were obtained for 318 (100%) and 304 (96%) isolates respectively. Isolates not typable according to the NG-STAR scheme (n=14) carried heterogeneous 23S rRNA alleles. Most prevalent NG-MAST and NG-STAR types in each MLST cluster are shown in Table S3.

Isolates belonging to MLST1901 were significantly overrepresented in 2014-2016 (15/19) and none of those were isolated after 2018. Also, MLST7363 isolates were significantly overrepresented in 2014-2016 (15/17). Remarkably, MLST7827 isolates were significantly overrepresented in 2017-2019 (50/63) of which 41/50 (82%) were isolated in 2018-2019 (Table S2). These results indicate the recent emergence of MLST7827 which became the dominant reduced susceptible strain in Amsterdam during 2017-2019, instead of the previously dominating MLST1901 and MLST7363 strains.

#### Azithromycin

Azithromycin resistance was found for 15/318 (5%) isolates and resistance was significantly associated with MLST9363 (Table S2). Remarkably, only one isolate carried one 23S rRNA allele with the known resistance mutation C2611T while all other isolates carried wild-type 23S rRNA genes. A mosaic *mtrR* gene was identified in 13/15 (87%) resistant isolates of which 69% (9/13) belonged to MLST9363. The same mosaic *mtrR* gene was found only once in a susceptible isolate, belonging to MLST7363. The other two resistant isolates carried either a 35A deletion or a A39T mutation in the *mtrR* promoter region, but these mutations were also highly prevalent among susceptible isolates (Table 2).

**Table 2.**
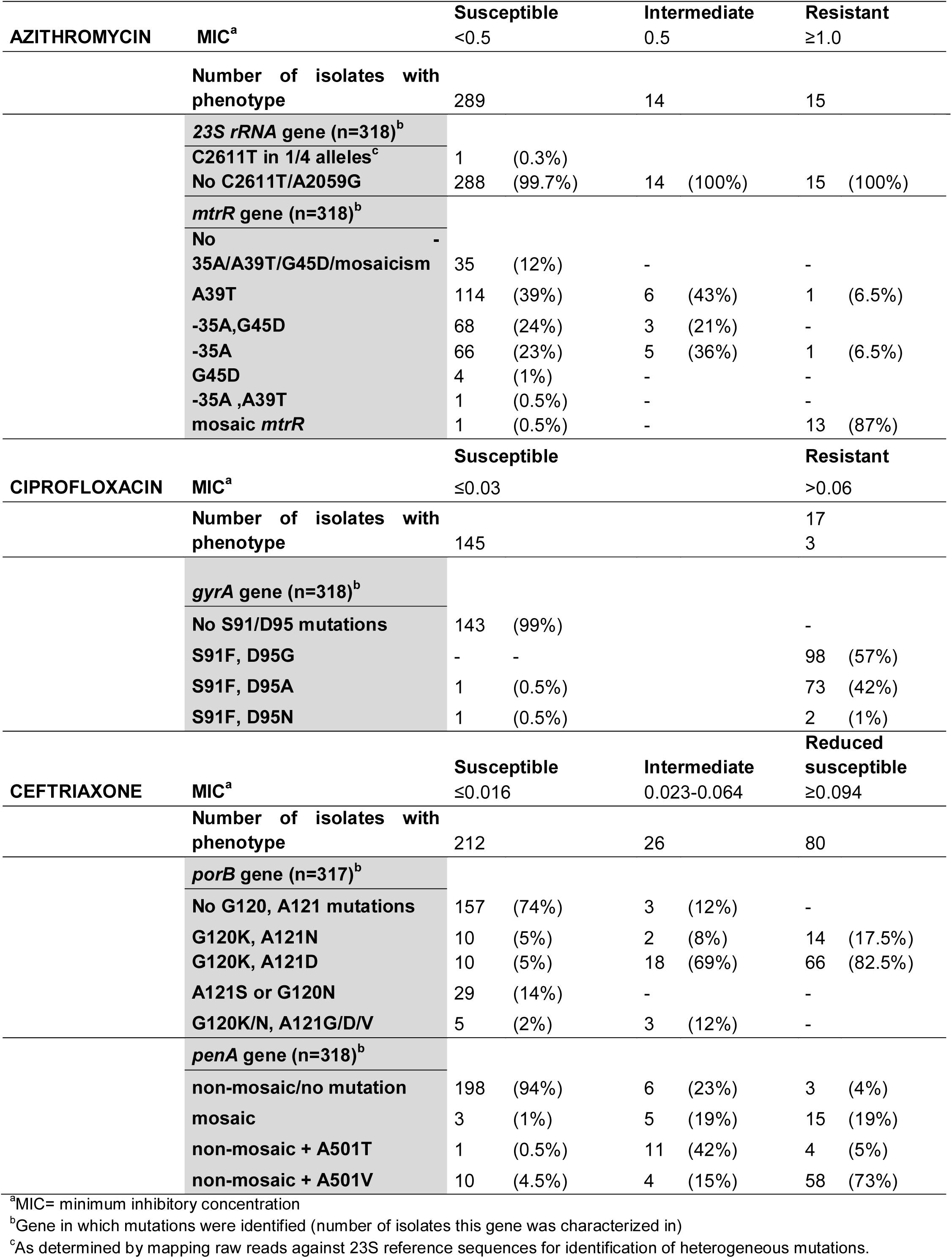
Phenotypic characterization versus identified resistance mutations.

#### Ciprofloxacin

Ciprofloxacin resistance was found for 173/318 (54%) isolates and significant associations were found with MLST1583, MLST1901, MLST7363 and MLST7827. MLST1599, MLST8156, MLST11428, MLST11864 and MLST11990 were significantly associated with susceptibility (Table S2). All resistant isolates carried the *gyrA* S91F mutation (173/173) which was found in only 2/145 (1%) susceptible isolates, showing the important role of this mutation in ciprofloxacin resistance. All isolates with an additional D95G mutation were resistant, whereas the 2 susceptible isolates carried either the D95A or N mutation (Table 2).

#### Ceftriaxone

As a result of the selection strategy of this study, 80/318 (25%) isolates were ceftriaxone reduced susceptible and 26/318 (8%) were intermediate susceptible. Intermediate- and reduced susceptibility was significantly associated with MLST1901, MLST7827 and MLST7363, and susceptibility with MLST1599, MLST8156 and MLST9363 (Table S2). The majority of susceptible isolates carried a *porB* gene without G120/A121 mutations (74%) and a non-mosaic *penA* gene without an A501 mutation (94%). Co-occurence of *porB* G120K/A121D mutations was found in 82.5% of the reduced susceptible-, 69% of the intermediate susceptible- and 5% of the susceptible isolates, indicating an important role of these mutations in the resistance mechanism. *PenA* mosaicism was found in 15/80 (19%) reduced susceptible isolates which all belonged to MLST1901. Notably, the mosaic *penA* allele was also found in intermediate susceptible MLST1901-(3/4) and MLST7363 isolates (2/11). The non-mosaic *penA* gene with A501T mutation was found in the majority of intermediate susceptible isolates (42%) of which 82% (9/11) belonged to MLST7363. The majority of reduced susceptible isolates carried a non-mosaic *penA* gene with A501V mutation (73%) of which 97% (56/58) belonged to MLST7827 (Table 2). The 4 susceptible MLST7827 isolates carried either a single- or no *porB* mutation and 2/4 also carried a non-mosaic *penA* without an A501 mutation. *PorB* G120K/A121D mutations that were found outside the MLST7827 cluster did not influence ceftriaxone susceptibility when co-occurring with a non-mosaic *penA* gene without mutations, except for 3 isolates in the MLST10314 cluster (Figure 1). However, intermediate- or reduced susceptibility was observed outside the MLST7827 cluster when both *porB* mutations and either *penA* mosaicism or A501 mutations co-occurred (e.g. in MLST1901 and MLST7363 clusters). Moreover, isolates carrying a *penA* A501V mutation but lacking the *porB* mutations were susceptible (e.g. in MLST1583 cluster). These findings indicate that ceftriaxone resistance is strongly influenced by an interplay between mutations in *penA* and *porB* and that some of these mutations show a stronger effect on MIC than others, thus gradually influencing the susceptibility.

### Close genetic relatedness between Dutch- and other European MLST7827 strains

The genetic relatedness of all 224 Dutch- and global MLST7827 isolates publicly available in the pubMLST database was assessed. Available metadata showed that a large proportion of the global MLST7827 isolates were from 2011-2013 (43%) and from Asia (42%). The Dutch isolates from this study accounted for 28% of the MLST7827 isolates in the database. Regarding available phenotypic data, 100% (192/193) of the isolates were ciprofloxacin resistant but only 3% (3/119) were azithromycin resistant. Regarding ceftriaxone, 36% (70/195) were intermediate- and 47% (92/195) were reduced susceptible (Table 3).

**Table 3.** Metadata available for *Neisseria gonorrhoeae* isolates belonging to Multi-Locus Sequence Type 7827 obtained from the pubMLST database in August 2020.

| Number of MLST7827 isolates in pubMLST database |  | 224 |
| --- | --- | --- |
| **Country/continent of isolation** | Netherlands | 63 (28%) |
|  | Norway | 30 (13%) |
|  | Other European countries +UK | 22 (10%) |
|  | America (continent) | 13 (6%) |
|  | Asia | 94 (42%) |
|  | New-Zealand | 2 (1%) |
| **Year of isolation** | 2011-2013 | 96 (43%) |
|  | 2014-2016 | 45 (20%) |
|  | 2017-2019 | 79 (35%) |
|  | NA<sup>a</sup> | 4 (2%) |
| **Ciprofloxacin MIC<sup>b</sup>** | ≤0.03 | 1 |
|  | >0.06 | 192 |
|  | NA<sup>a</sup> | 31 |
| **Azithromycin MIC<sup>b</sup>** | <0.5 | 101 |
|  | 0.5 | 15 |
|  | ≥1.0 | 3 |
|  | NA<sup>a</sup> | 105 |
| **Ceftriaxone MIC<sup>b</sup>** | ≤0.016 | 33 |
|  | 0.023-0.08 | 70 |
|  | ≥0.094 | 92 |
|  | NA<sup>a</sup> | 29 |

<sup>a</sup>NA= data not available in pubMLST database
<sup>b</sup>MIC=minimum inhibitory concentration

Alignment of the MLST7827 core genome resulted in an alignment length of 2,166,113 sites. Recombination filtered variants alignment resulted in an alignment length of 7,526 sites, on which the phylogenetic tree was based (Figure 2). The midpoint-rooted phylogenetic tree showed two main lineages: lineage A with mainly Asian isolates from 2011-2013, and lineage B with 3 distinct clusters. Cluster 1 and 3 mainly contained European isolates from 2014-2019 and cluster 2 mainly contained Asian, but also American and European isolates from 2011-2013. Dutch isolates from 2017-2019 were only found in cluster 1, together with Norwegian- and other European isolates and one American isolate. The median SNP distances within clusters 1, 2 and 3 were 34, 89 and 33 respectively. This indicated stronger genetic relatedness among European isolates in clusters 1 and 3 than among isolates in cluster 2, which were mainly Asian. Cluster 1 contained most ceftriaxone reduced susceptible isolates carrying non-mosaic *penA* alleles with A501V mutation and *porB* G120K/A121D mutations. In contrast, lineage A mainly contained susceptible isolates carrying *penA* and *porB* alleles without mosaicism or mutations. Isolates in cluster 2 and 3 mainly carried the non-mosaic *penA* with A501V mutations, however a variety of *porB* alleles was found in those clusters. Overall, ceftriaxone reduced susceptibility was associated with the co-presence of the *penA* A501V and *porB* G120/A121 mutations among global MLST7827 isolates.

**Figure 2.**
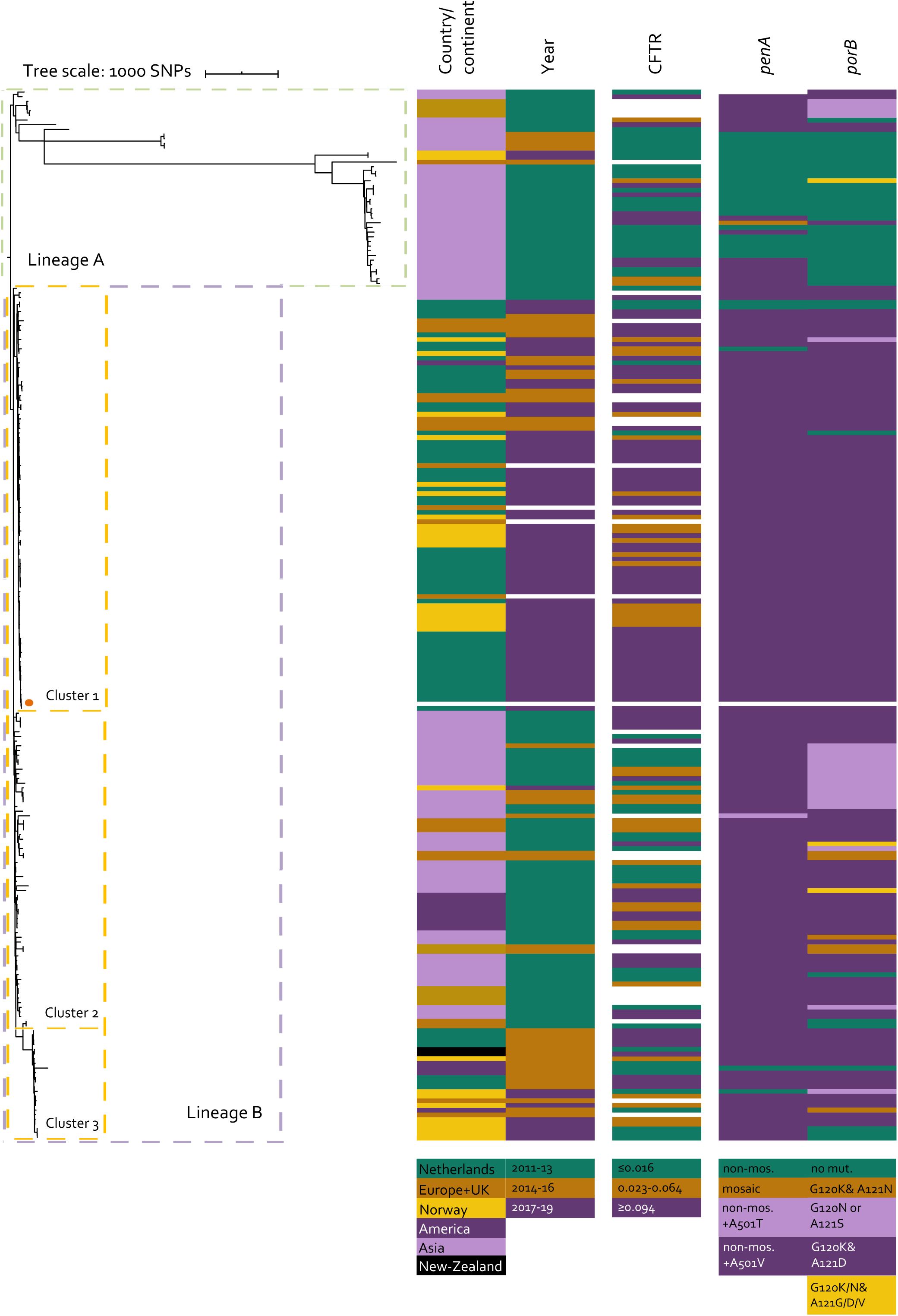
Recombination filtered midpoint-rooted phylogenetic tree based on core genome SNPs including all 224 publicly available global *Neisseria gonorrhoeae* isolates belonging to Multi-Locus Sequence Type 7827. A DutchMLST7827 isolate was used as reference strain and its branch is visualized with an orange dot. Metadata includes: country/continent; year of isolation; ceftriaxone (CFTR) Minimum Inhibitory Concentrations visualized as susceptible (green), intermediate susceptible (orange) and reduced susceptible (purple); types of *penA* and *porB* genes. White bars indicate missing data. Dashed lined boxes define separate lineages and clusters.

## Discussion

This genomic population study extends the NG-MAST and *penA* typing study on *Ng* isolates from Amsterdam, isolated up until 2017 (14). Back then, a shift in *penA* (from mosaic to non-mosaic with an A501T or A501V mutation) and NG-MAST was found among ceftriaxone reduced susceptible isolates. In the current study, this phenomenon was examined in more detail adding more recent isolates and using whole genome sequencing. The results showed that previous observations represented a shift from MLST1901 to MLST7363, and more recently to MLST7827. The emergence of the MLST7827 strain with reduced susceptibility to cephalosporins and resistance to ciprofloxacin in Amsterdam is in line with previously published surveillance articles from different parts of the world. Peng et al. reported MLST7827 as the most prevalent MLST in China already in 2012-2013, although then this MLST was not particularly associated with cephalosporin reduced susceptibility (31). When this strain emerged in Norway in 2016-2018 it was associated with cephalosporin reduced susceptibility (13) and we now confirm that the same emergence has occurred in the Netherlands during the last 3 years. Since isolates from other European countries from the last few years are scarce, these results can only suggest the circulation of this ST in other parts of Europe.

Among the selection of Dutch isolates, the co-occurrence of a non-mosaic *penA* allele with A501V mutation and *porB* G120K/A121D mutations was determinative for ceftriaxone reduced susceptibility. Remarkably, those mutations were already found in cephalosporin reduced susceptible isolates from Korea from 2001-2007, although these isolates belonged to different NG-MASTs (32). From 2007 onwards, cephalosporin resistance was often associated with MLST1901 and/or a mosaic *penA* allele, although other MLST and *penA* types were found among ceftriaxone reduced susceptible isolates in the last decade as well. In China, *Ng* isolates belonging to MLST7363 and carrying a mosaic *penA* were responsible for reduced susceptibility in 2012-2013 (31). We found both MLST1901 and MLST7363 to be associated with reduced susceptibility from 2014-2016 in Amsterdam as well, confirming that these strains were dominating the circulating reduced susceptible strains worldwide in previous years. The mosaic *penA* was found in only a small subset of the MLST7363 isolates, and these were genetically very distinct from MLST7363 isolates with a non-mosaic *penA* allele with A501T mutation (Figure 1). In 2017, Abrams et al. reported a reduced susceptible isolate that belonged to a different ST and lacked the mosaic *penA* allele (33). Additional to the shift towards other non-mosaic *penA* alleles found previously in Amsterdam, we here identified the association between reduced susceptibility and MLST7827 isolates carrying *porB* G120K/A121D mutations. Among the global MLST7827 isolates, both *penA* and *porB* mutations were determinative for ceftriaxone reduced susceptibility. Altogether these findings indicate that the cephalosporin resistance mechanism is not determined by a single genetic variation but that different routes towards resistance exist. This multifactorial nature also suggests additional and yet unresolved genetic variations involved in resistance.

Although this study included only a few azithromycin resistant isolates, the results suggest a replacement of mutations in 23S rRNA by mosaic *mtrR* as main determinant for azithromycin resistance. Previous research on Dutch isolates from 2008-2015 showed a high prevalence of 23S rRNA mutations among azithromycin resistant isolates with variable genetic backgrounds (34). In contrast, mosaic *mtrR* genes instead of mutations in 23S rRNA were found among azithromycin resistant isolates included in this study. Further research on larger numbers of azithromycin resistant isolates is needed to confirm this observation.

Phylogenetic analysis including the 318 Dutch isolates revealed two separate lineages. Isolates in lineage B were significantly associated with bi- or homosexual intercourse and resistance or reduced susceptibility against azithromycin, ciprofloxacin and ceftriaxone was significantly overrepresented as well. Previous studies state that MSM are more often infected with isolates that are reduced susceptible to antibiotics, probably because of higher prevalence of bacterial STIs in MSM. This leads to higher antibiotic exposure and increases selection pressure for antimicrobial resistance (35)(36).

Importantly, the percentage of ceftriaxone reduced susceptibility found in this study is not representative for the *Ng* population found in STI clinic visitors in Amsterdam, since isolates were selected for WGS based on ceftriaxone susceptibility. Routine monitoring of ceftriaxone susceptibility showed that only 82/7323 (1.1%) isolates were reduced susceptible during the study period. Although ceftriaxone resistant isolates have not yet been found in the Netherlands, the emergence of MLST7827 isolates raises the question if these isolates will evolve towards resistant isolates in the future. Until now, reported resistant isolates always carried a mosaic *penA* gene which was found in only one global MLST7827 isolate. So far, the *penA* A501V and *porB* G120/A121 mutations are only associated with reduced susceptibility to ceftriaxone. High recombination rates in *Ng* enable the exchange of resistance mutations in other genes, which could cause a further reduction in susceptibility and ultimately lead to resistance. Identification of the MLST7827 strain in multiple European countries over the last years shows its ability to spread quickly, which underlines the need for global surveillance to track the prevalence and development of this strain in the future.

### Ethical declaration

According to the Dutch Medical Research Act Involving Human Subjects, no additional ethical approval was required for this study (W20_451 # 20.498).

## Acknowledgements

The authors would like to thank Boas van der Putten for his valuable input and careful reading of the manuscript.

