## Supplementary material for "Emergence of a cephalosporin reduced susceptible Neisseria gonorrhoeae clone between 2014-2019 in Amsterdam, the Netherlands, revealed by a genomic population analysis"

**Table S1. Significance of associations between lineage A (n=102 isolates) and lineage B (n=216 isolates) and patient characteristics, tested with Chi-square tests.**

|  |  | **Prevalence in lineage A** | **Prevalence in lineage B** | **p-value** |
| --- | --- | --- | --- | --- |
| **Age^a^** | <24 | 33% | 12% |  |
|  | 24-34 | 42% | 55% |  |
|  | ≥35 | 25% | 33% | **2.74E-05** |
| **Sex** | Male | 78% | 97% | **1.53E-07** |
| **Sexual preference** | MSM or bisexual | 66% | 95% | **6.57E-12** |
| **Year of isolation** | 2017-2019 | 47% | 66% | **0.002** |
| **AZI^b^** | Resistant | 1% | 6% | **0.04** |
| **CIP** | Resistant | 39% | 61% | **0.0003** |
| **CFTR** | Reduced susceptible | 5% | 35% | **2.38E-08** |

^a^Groups defined for each characteristic as follows: Age: 1) <24, 2) 24-34 and 3) ≥35 years; Sexual preference: 1) exclusively reported heterosexual intercourse and 2) reported bi- or homosexual intercourse; Year of isolation: 1)2014-2016 and 2) 2017-2019; Phenotype: 1) Isolates with MIC assigned as susceptible or intermediate susceptible 2) Isolates with MICs assigned as resistant or reduced susceptible (in case of ceftriaxone).

^b^Fisher’s exact test performed because one of the groups contained <5 isolates

**Table S2. Significance of associations between main MLST clusters and patient- or strain characteristics, tested with Fisher’s exact tests with Bonferroni correction for multiple testing.**

|  |  | **Year of isolation^a^** | | **Sex** |  | **Age** |  | **Sexual preference** | | **AZI** | |  | | **CIP** |  | **CEFTR** | |
| --- | --- | --- | --- | --- | --- | --- | --- | --- | --- | --- | --- | --- | --- | --- | --- | --- | --- |
|  |  | 2017-2019 | | Male |  | ≤30 years |  | MSM or bisexual | | Resistant | | | | Resistant |  | Reduced susceptibility | |
| **MLST cluster** | **Isolates in cluster** | Prevalence | p-value | Prevalence | p-value | Prevalence | p-value | Prevalence | p-value | Prevalence | p-value | | | Prevalence | p-value | Prevalence | p-value |
| **1583** | 13 | **100%** | **0.039** | 100% | 1 | 46% | 1 | 92% | 1 | 0% | 1 | | | **100%** | **0.006** | 0% | 0.104 |
| **1588** | 5 | 80% | 1 | 80% | 1 | 80% | 1 | 60% | 1 | 0% | 1 | | | 80% | 1 | 0% | 1 |
| **1599** | 17 | 59% | 1 | 100% | 1 | 47% | 1 | 100% | 1 | 0% | 1 | | | **0%** | **1.56E-05** | **0%** | **0.019** |
| **1901** | 19 | **21%** | **0.009** | 95% | 1 | 79% | 0.88 | 89% | 1 | 5% | 1 | | | **100%** | **1.87E-04** | **79%** | **5.13E-09** |
| **7363** | 17 | **12%** | **6.79E-04** | 71% | 0.198 | 35% | 1 | 71% | 1 | 6% | 1 | | | **94%** | **0.011** | **24%** | **4.55E-05** |
| **7822** | 12 | 75% | 1 | 92% | 1 | 58% | 1 | 92% | 1 | 0% | 1 | | | 92% | 0.14 | 0% | 1 |
| **7827** | 63 | **79%** | **0.009** | 98% | 0.421 | 43% | 1 | 94% | 0.292 | 0% | 1 | | | **100%** | **9.7E-19** | **89%** | **2.81E-30** |
| **8135** | 5 | 100% | 1 | 40% | 0.099 | 100% | 1 | **0%** | **5.83E-04** | 0% | 1 | | | 0% | 0.335 | 0% | 1 |
| **8143** | 8 | 38% | 1 | 100% | 1 | 38% | 1 | 100% | 1 | 0% | 1 | | | 100% | 0.161 | 0% | 0.1 |
| **8156** | 28 | 64% | 1 | 96% | 1 | 39% | 1 | 100% | 1 | 0% | 1 | | | **0%** | **9.4E-10** | **0%** | **1.59E-04** |
| **8163** | 7 | 57% | 1 | 57% | 0.308 | 100% | 0.286 | **0%** | **7.94E-06** | 0% | 1 | | | 0% | 0.066 | 0% | 1 |
| **9363** | 18 | 61% | 1 | 100% | 1 | 33% | 1 | 100% | 1 | **50%** | **0.0042** | | | 50% | 1 | **0%** | **0.011** |
| **10314** | 9 | 100% | 0.229 | 100% | 1 | 33% | 1 | 100% | 1 | 0% | 1 | | | 100% | 0.083 | 0% | 1 |
| **11428** | 12 | 33% | 1 | 100% | 1 | 50% | 1 | 100% | 1 | 0% | 1 | | | **0%** | **0.001** | 0% | 0.182 |
| **11864** | 11 | 55% | 1 | 100% | 1 | 45% | 0.072 | 100% | 1 | 0% | 1 | | | **0%** | **0.002** | 0% | 0.331 |
| **11990** | 9 | 33% | 1 | **22%** | **1.14E-05** | 100% | 1 | **0%** | **9.74E-08** | 0% | 1 | | | **0%** | **0.013** | 0% | 0.572 |
| **13292** | 5 | 80% | 1 | 100% | 1 | 20% | 1 | 100% | 1 | 0% | 1 | | | 0% | 0.335 | 0% | 1 |
| **<5 iso-lates^b^** | 59 | 53% | 1 | 92% | 1 | 64% | 1 | 86% | 1 | 93% | 1 | | | 41% | 0.367 | **8%** | **0.004** |
| **Total** | **317^c,d^** |  |  |  |  |  |  |  |  |  | | |  |  |  |  |  |

^a^Groups defined for each characteristic as follows: Year of isolation: 1)2014-2016 and 2) 2017-2019; Age:1) ≤30 years and 2) >30 years; Sexual preference: 1) exclusively reported heterosexual intercourse and 2) reported bi- or homosexual intercourse; Phenotype: 1) Isolates with MIC assigned as susceptible or intermediate susceptible 2) Isolates with MICs assigned as resistant or reduced susceptible (in case of ceftriaxone).

^b^MLST clusters which included <5 isolates were taken together

^c^One strain could not be typed according to the MLST typing scheme since one locus was only partially assembled

^d^Duplicate patients were added only once when tested for associations with patient characteristics (n=314 patients)

**Table S3. Most prevalent NG-MAST and NG-STAR types found in each main MLST cluster.**

| **MLST cluster** | **Number of isolates** | **Main NG MAST type** | **Main NG STAR type** |
| --- | --- | --- | --- |
| 1583 | 13 | 15589 (92%) | 1340 (54%) |
| 1588 | 5 | NA^a^ | NA |
| 1599 | 17 | 11461 (65%) | 520 (76%) |
| 1901 | 19 | 1407 (58%) | 90 (32%) |
| 7363 | 17 | 2400 (35%) | 158 (53%) |
| 7822 | 12 | 14994 (33%) | 1387 (42%) |
| 7827 | 63 | 10368 (35%) / 2318 (33%) | 38 (89%) |
| 8135 | 5 | 387 (60%) | 729 (60%) |
| 8143 | 8 | 5624 (50%) | 426 (50%) |
| 8156 | 28 | 5441 (71%) | 442 (89%) |
| 8163 | 7 | 2 (29%) | 84 (86%) |
| 9363 | 18 | 12302 (28%) | 168 (28%) |
| 10314 | 9 | NA | 1387 (44%) |
| 11428 | 12 | 2992 (58%) | 63 (92%) |
| 11864 | 11 | 18234 (27%) | 439 (91%) |
| 11990 | 9 | 14376 (56%) | 962 (56%) |
| 13292 | 5 | 9208 (80%) | 439 (80%) |

^a^In case of ‘NA‘ there is no dominant ST found in that MLST cluster.
